# Differences in Early and Late Pattern-Onset Visual-Evoked Potentials between Self-Reported Migraineurs and Controls

**DOI:** 10.1101/733816

**Authors:** Chun Yuen Fong, Wai Him Crystal Law, Jason Braithwaite, Ali Mazaheri

## Abstract

Striped patterns have been shown to induce strong visual illusions and discomforts to migraineurs in the literature. Previous research has suggested that those unusual visual symptoms be linked with the hyperactivity on the visual cortex of migraine sufferers. The present study searched for evidence supporting this hypothesis by comparing the visual evoked potentials (VEPs) elicited by striped patterns of specific spatial frequencies (0.5, 3, and 13 cycles) between a group of 29 migraineurs (17 with aura/12 without) and 31 non-migraineurs. In addition, VEPs to the same stripped patterns were compared between non-migraineurs who were classified as hyperexcitable versus non-hyperexcitable using a previously established behavioural pattern glare task. We found that the migraineurs had a significantly increased N2 amplitude for stimuli with 13 cpd gratings but an attenuated late negativity (LN: 400 – 500 ms after the stimuli onset) for all the spatial frequencies. Interestingly, non-migraineurs who scored as hyperexcitable appeared to have similar response patterns. We proposed that the enhanced N2 could reflect disruption of the balance between parvocellular and magnocellular pathway, which is in support of a grating-induced cortical hyperexcitation mechanism on migraineurs. On the other hand, the attenuation of the late negativity could reflect a top-down feedback mechanism to suppress visual processing of an aversive stimulus.

## 1. Introduction

Research on visual stress has proposed that the aversive visual responses to gratings of specific spatial frequencies could be due to the cortical hyperexcitation of the visual cortex (Evans, Cook, Richards, & Drasdo, 1994; Harle, Shepherd, & Evans, 2006; Wilkins, 1995). This hypothesis was mainly based on the research of migraine patients, who have shown evidence of hypersensitivity to gratings in various behavioural and neurophysiological studies (Aurora & Wilkinson, 2007).

Visual evoked potentials (VEPs) are EEG responses that are both time and phase-locked to the onset of a visual stimulus. Studies using magnetoencephalography (MEG) and functional magnetic resonance imaging (fMRI) revealed that the initial observable VEP components, C1 and N75 are originated from the V1 (Di Russo et al., 2005; Foxe et al., 2008; Hatanaka et al., 1997). On the other hand, the generators of the later components, P100 and N145 have been localised to the extrastriate visual cortex (V2 – V4; Di Russo et al., 2005; Lehmann, Darcey, & Skrandies, 1982; Schroeder et al., 1995; Vanni et al., 2004). Importantly, the peak amplitude and latency of these early components are consistently found to be influenced by the psychophysical features of the visual stimuli such as spatial frequency, contrast and colour (Ellemberg, Hammarrenger, Lepore, Roy, & Guillemot, 2001; Oelkers et al., 1999; Souza et al., 2008).

The difference in VEP peak amplitude or latency obtained from group comparisons (clinical vs control) may indicate impairment of early visual processing in the testing patient groups (e.g. migraine; Afra, Cecchini, DePasqua, Albert, & Schoenen, 1998; Shibata, Osawa, & Iwata, 1997,1998). For example, prolonged or increased P100 was found to be associated with the visual hallucinations and other forms of visual disturbances amongst the Parkinson and schizophrenic populations. Among all the clinical populations who are highly susceptible to visual disturbances, the VEPs of migraine sufferers (with/without aura) have been most widely studied (e.g. Di Russo et al., 2005; Shibata, Yamane, Iwata, & Ohkawa, 2005). Previous studies have found that migraine sufferers have increased peak-to-peak amplitudes for both N75-P100 and P100-N145 interictally compared to healthy controls (Oelkers et al., 1999; Shibata, Osawa, & Iwata, 1997, 1998). Heterogenous measurements on peak latencies have also been reported, with studies revealing a rather mixed set of findings including prolonged, shortened or unchanged peak latencies in migraineurs (Afra et al., 1998; Oelkers et al., 1999; Shibata, Osawa, & Iwata, 1997; Tagliati, Sabbadini, Bernardi, & Silvestrini, 1995). One particular study, Oelkers et al. (1999), found a prolonged N2 latency in migraine patients compared to control (Oelkers et al., 1999; Yilmaz et al., 2001). They attempt to account for this finding by proposing that the N2 component consists of a parvocellular “N130” (contour processing) and magnocellular “N180” complex (luminance processing). The prolonged N2 latency in the migraineurs could emerge due to an enhancement in the N180 as a result of the imbalance of the two visual pathways.

In addition to the early VEP components, previous research has also found that migraineurs also exhibit abnormality for late components (e.g. Drake, Pakalnis, & Padamadan, 1989; Mazzotta, Alberti, Santucci, & Gallai, 1995; Puca & Tommaso, 1999). The P3, is a positive potential, peaking maximally over central-parietal electrodes at around 300ms post-stimulus after ‘oddball stimuli’, infrequent deviant stimuli (e.g. an X) occurring within a train of frequent standard stimuli. The latency of the P300 has been suggested to reflect the time it took participants to discriminate/categorise the oddball stimulus as deviant, while the amplitude decrease with their confidence in that discrimination (see Picton, 1992 for review). Previous research has found that migraineurs often have a longer P3 latency in oddball paradigm (Bockowski et al., 2004; Chen et al., 2007; Drake et al., 1989; Mazzotta et al., 1995) indicating a more prolonged time needed to discriminate the target stimuli. Studies with the primary focus on VEP between 400 – 700 ms have been somewhat limited and often reported with contradictory findings (Mickleborough et al., 2013; Tommaso et al., 2009; Steppacher, Schindler, & Kissler, 2016). This does suggest though that the differences in the VEPs of migraineurs and non-migraineurs are not restricted to early exogenous ERPs (i.e. occurring within 200ms post-stimulus onset).

### 1.1. Current study

Migraineurs are known to be more susceptible to visual discomforts and distortions in viewing gratings of 2 – 4 cycles per degree (cpd), which is also known as pattern glare (Evans & Stevenson, 2008). Here we set out to compare the VEPs elicited by gratings having 3 different spatial frequencies (0.5, 3, and 13 cycles per degree: cpd) between migraineurs and non-migraineurs, with N2 as our primary focus. Evans and Stevenson (2008) proposed that pattern glare is a consequence of over-stimulation within the same nerve network on a hyperexcitable visual cortical area. Therefore, in parallel with migraineurs, non-clinical populations who have hyperexcitable visual cortex might also show symptoms of pattern glare. In order to investigate this possibility, we compared the VEPs elicited by gratings between a larger group of non-migraineurs separated into a hyperexcitable and non-hyperexcitable group according to the behavioural responses to a pattern glare task.

## 2. Methods

### 2.1. Participants

Twenty-nine migraine female patients (mean age = 20.9) and 31 healthy female controls (mean age = 19.4) participated in the first study. Seven additional healthy female subjects were later added to the second part of the study comparing hyperexcitable and non-hyperexcitable non-migraineurs (total: 38, mean age = 19.3). All the participants had normal/corrected visual acuity (20/25 or better). The participants in the control group reported no history of migraine nor any other neurological and psychiatric conditions. In the migraine group, 17 were classified as having migraine with aura and 12 with migraine without aura based on the criteria proposed by the International Headache Society (Olesen, 2018). They were not regularly taking prophylactic medications (and had not taken one within 2 weeks of the EEG session), nor had chronic migraine, motor migraine aura symptoms or any other forms of neurological or psychiatric conditions. Finally, these participants were studied interictally, such that they did not have a migraine attack in the week leading up to the EEG recordings, and followed up for at least 2 weeks after the recordings. This study has been approved by the Ethics Committees of the University of Birmingham.

### 2.2. Stimuli, Apparatus and Questionnaires

The stimuli used in this experiment included 3 square-wave achromatic gratings: a low-frequency grating (LF) of 0.5cpd, a medium-frequency grating of 3cpd, and a high-frequency grating of 13cpd (see Figure 7 for an example of the grating). All stimuli were presented at the centre of a 20-inch Dell P2210 LCD computer screen (60Hz refresh rate and 1680×1050 pixels screen resolution) using E-prime v2.0 software, with a background luminance of 20 cd/m^2^. The Michelson contrast of all the 3 gratings was 0.70 (cd/m^2^). Each of them had an identical shape with the maximum height x width of 140 mm x 180 mm with the shape of a mild ellipse different in spatial frequency (cycles-per-degree: cpd). The viewing distance was fixed at 80 cm, which gave a visual angle of 9.93 x 12.68 degrees.

Participants also completed 2 questionnaires which measure the trait-based predisposition to anomalous perceptions: the *Cortical Hyperexcitability index – II (CHi-II)* (Fong, Takahashi, & Braithwaite, 2019) and the *Cardiff Anomalous Perceptual Scale (CAPS)* (Bell, Halligan, & Ellis, 2005). The *CHi-II* has three factors representing different types of anomalous visual experiences, namely, (i) *Heightened Visual Sensitivity and Discomfort (HVSD)*; (ii) *Aura-like Hallucinatory Experiences (AHE)*; (iii) *Distorted Visual Perception (DVP)*. Similar to *CHi-II, CAPS* could be broken down into 3 components: *Temporal Lobe Experience (TLE), Chemosensation (CS) and Clinical Psychosis (CP)*.

### 2.3. Procedures

#### 2.3.1. The Pattern-glare task

The participants first completed a computerised version of the pattern-glare task, where observers reported phantom visual distortions from viewing highly irritable visual patterns (PG; Braithwaite, Mevorach, & Takahashi, 2015; Fong et al., 2019; Evans & Stevenson, 2008). Each trial began with a 12-second-presentation of one of the three gratings (presented in a pseudo-random order; see Figure 1 for the grating). Participants were instructed to gaze on a fixation point locating at the centre of the grating.

**Figure 1.**
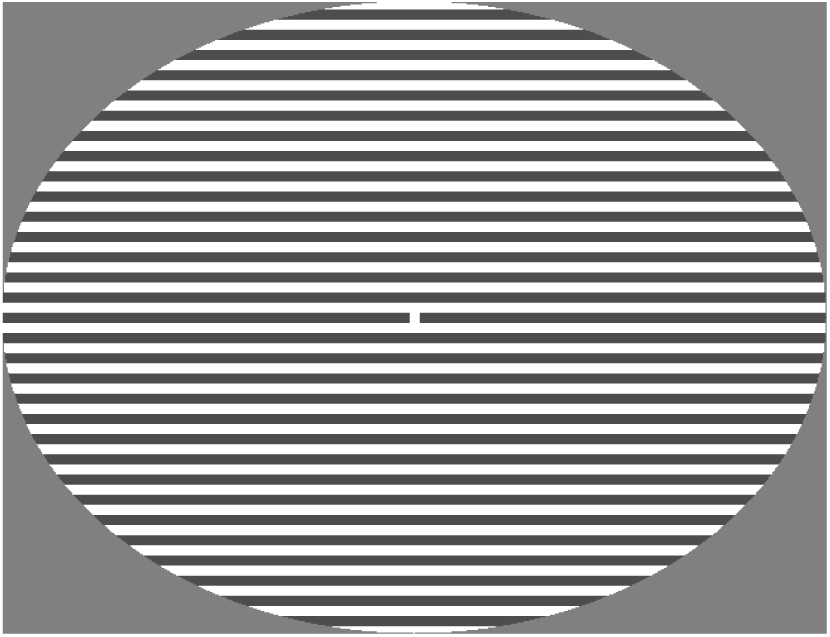
An example of the medium frequency square-wave grating

After the stimuli presentation, participants gave their responses on the intensity/ strength of the associated visual distortions (AVD; visual pain, physical eye strain, unease, nausea, headache, dizziness, light-headedness, faint, shadowy shape, illusory stripes, shimmering, flickering, jitter, zooming, blur, bending of lines, and colour distortions: red, green, blue, yellow) they had experienced using a 7-point Likert scale (0 = not at all, 6 = extremely; see Figure 2 for the trial sequence). The responses for each AVD were added together to give a total AVD score for that grating (range = 0 – 120). AVD scores for each grating were obtained from the average AVD of the 3 repetitions for that spatial frequency.

**Figure 2.**
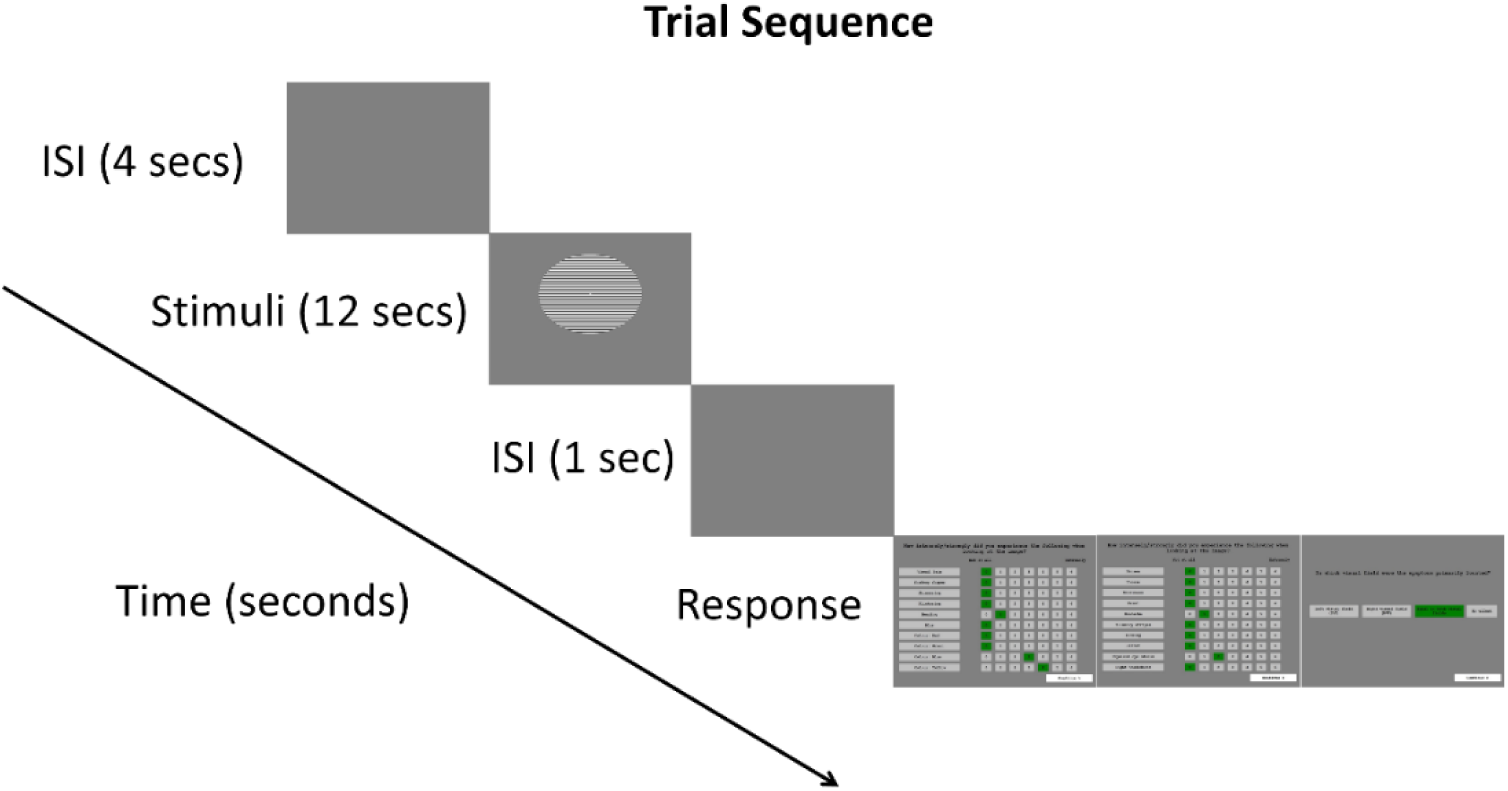
Trial sequence for the PG task

#### 2.3.2. EEG task

After the completion of the pattern glare task, we examined the EEG responses of participants elicited by the presentations of the gratings. Each trial began with a 2s-fixation period where the participants were asked to fixate on a cross at the centre of the screen after which one of the gratings (HF, MF or LF) was then presented. Participants were instructed to keep their focus at the centre of the stripe patterned-disc. Then, they were asked to either hit the left-click by their index finger when their visual discomforts/distortions had reached the maximum (typically around 2 to 10 seconds) or the right-click by their middle finger when they did not experience any forms of visual discomforts/distortions at all after 8 seconds counting in their minds. Each trial was separated by an 8s inter-stimulus interval (see Figure 3 for the EEG task trial sequence).

**Figure 3.**
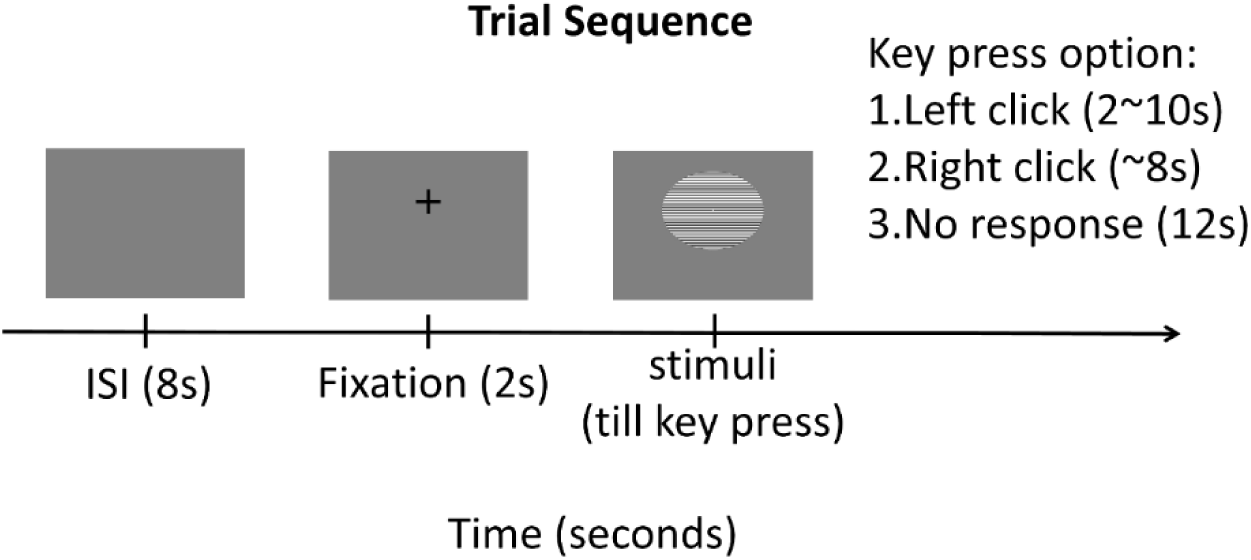
Trial sequence for the EEG task. The behavioural response (key press) was not analysed in the current study.

Each grating was presented 50 times in pseudo-random order. The total 150 trials were divided into 10 blocks which were separating by breaks (the durations were entirely controlled by the subjects).

##### 2.3.2.1. EEG Data Acquisition

The EEG signal was recorded by the EEGO Sports system (ANT Neuro) and Waveguard caps containing 64 Ag/AgCl electrodes (10/10 systems including left and right mastoids). Electrodes at CPz was used as reference while AFz was used as ground. After analogue to digital conversion, EEG data (sampling rate 500 Hz, impedances < 20 kΩ) was amplified with a high pass of 0.8 Hz and low-pass of 30 Hz. Two pairs of bipolar EOG electrodes were used to measure both horizontal and vertical eye movements. One was placed at the outer canthi of left and right eyes while another pair was placed at the left lid-cheek junction and above left eyebrow. Heartbeat data, which was not used in the present study, was measured by placing a pair of ECG electrodes on the left and right chest (grounded at left collar bone).

##### 2.3.2.2. Pre-processing

The EEG data was pre-processed in Matlab using EEGLAB functions (version 14.1.2b; Delorme & Makeig, 2004). First, the data was epoched from −500 to 1500 ms around the stimulus onset. Next, the extracted epochs were broken down into 30 components by Independent Component Analysis (ICA) using Principal Component Analysis (PCA) as the extracting method. Components that reflected muscle and ocular artefacts (e.g. eye blinks) were removed. Trials were then manually inspected visually, and those with muscle artefacts and noise not corrected by the ICA were removed. Spherical interpolation function in EEGLAB was used to fix the bad electrodes in the data (see Ferree, 2000). Finally, CPz reference montage was replaced by average reference prior to any further analysis.

### 2.4. Design and Analysis

The cleaned epoched data was further analysed by FieldTrip software package (Oostenveld, Fries, Maris, & Schoffelen, 2011) using both confirmatory and exploratory approaches. The trials epoched around the onset of each of the visual gratings were averaged to obtain VEPs. The Oz electrode was chosen to measure the early VEP components according to the latencies after the stimulus onset (Di Russo et al., 2005; Khalil, Legg, & Anderson, 2000). The peak latency range (N2 and other components) for each of the gratings was defined by visual inspection, on the grand-averaged ERPs, collapsed across all participants. A two-tailed independent samples t-test was performed to assess the statistical significance of differences in the mean peak amplitude and peak latency within these time-intervals of the interest between groups. In addition, Bayesian analyses were conducted by JASP version 0.8.0.1, using the default Cauchy prior width (0.707) (Love et al., 2015; Rouder, Morey, Speckman, & Province, 2012). The analysis provides relative evidence and probability on whether the data are more in favour of the alternative hypothesis (H_A_) (*BF*_*10*_ > 1.0) or the null-hypothesis (H_0_) (*BF*_*10*_ < 1.0). For example, a *BF*_*10*_ of 0.1 suggests that the H_0_ is 10 times more likely than the H_A_. In general, a *BF*_*10*_ close to 1 are not informative, while a *BF*_*10*_ > 3 or < 0.33 can be interpreted as moderate evidence in favour of the H_A_ or H_0_ respectively (Jarosz & Wiley, 2014).

As an exploratory approach, any amplitude differences within the time window 0 – 700 ms between-group were assessed by non-parametric cluster-based permutation analysis (Maris & Oostenveld, 2007). The adjacent spatiotemporal sample data was first clustered if they exceeded a threshold of *p* < 0.05 (cluster-alpha). The cluster with a Monte Carlo *p*-value smaller than 0.025 was identified as significant (simulated by 1000 partitions), thus, showing a significant group difference in amplitude.

## 3. Results

### 3.1. Migraine vs healthy control

#### 3.1.1. VEP Component Analysis

As mentioned earlier, the peak latency range for each of the gratings was defined by visual inspection, on the grand-averaged ERPs, collapsed across all participants, at the occipital Oz electrode. The key component – N2 was defined as the mean amplitude between 170 – 240 ms for HF, 140 – 180 ms for MF, and 150 – 185 ms for LF. The independent samples t-test showed that the migraineurs had a more negative N2 amplitude in HF conditions than control group (mean: −2.67 μV vs −1.21 μV), t (58) = 2.744, uncorrected *p* = 0.008, False Discovery Rate (FDR) corrected *p* = .024, *BF*_*10*_ = 5.59 (see Figure 4 & Table 1).

**Table 1.** Results of independent t-test and Bayes factor on N2 amplitudes between migraineurs and control with standard error in parenthesis.

| Mean amplitude ($\mu\text{V}$) | Migraine (n = 29) | Control (n = 31) | *t*-stat | uncorrected *p*-value | *FDR*-corrected *p*-value | *BF*<sub>10</sub> |
| --- | --- | --- | --- | --- | --- | --- |
| **HF** | **-2.67 (0.40)** | **-1.21 (0.35)** | **2.74** | **.008** | **.024** | **5.59** |
| MF | -0.28 (0.49) | 0.80 (0.66) | 1.30 | .20 | .20 | 0.53 |
| LF | -3.39 (0.47) | -2.23 (0.57) | 1.56 | .12 | .18 | 0.73 |

**Figure 4.**
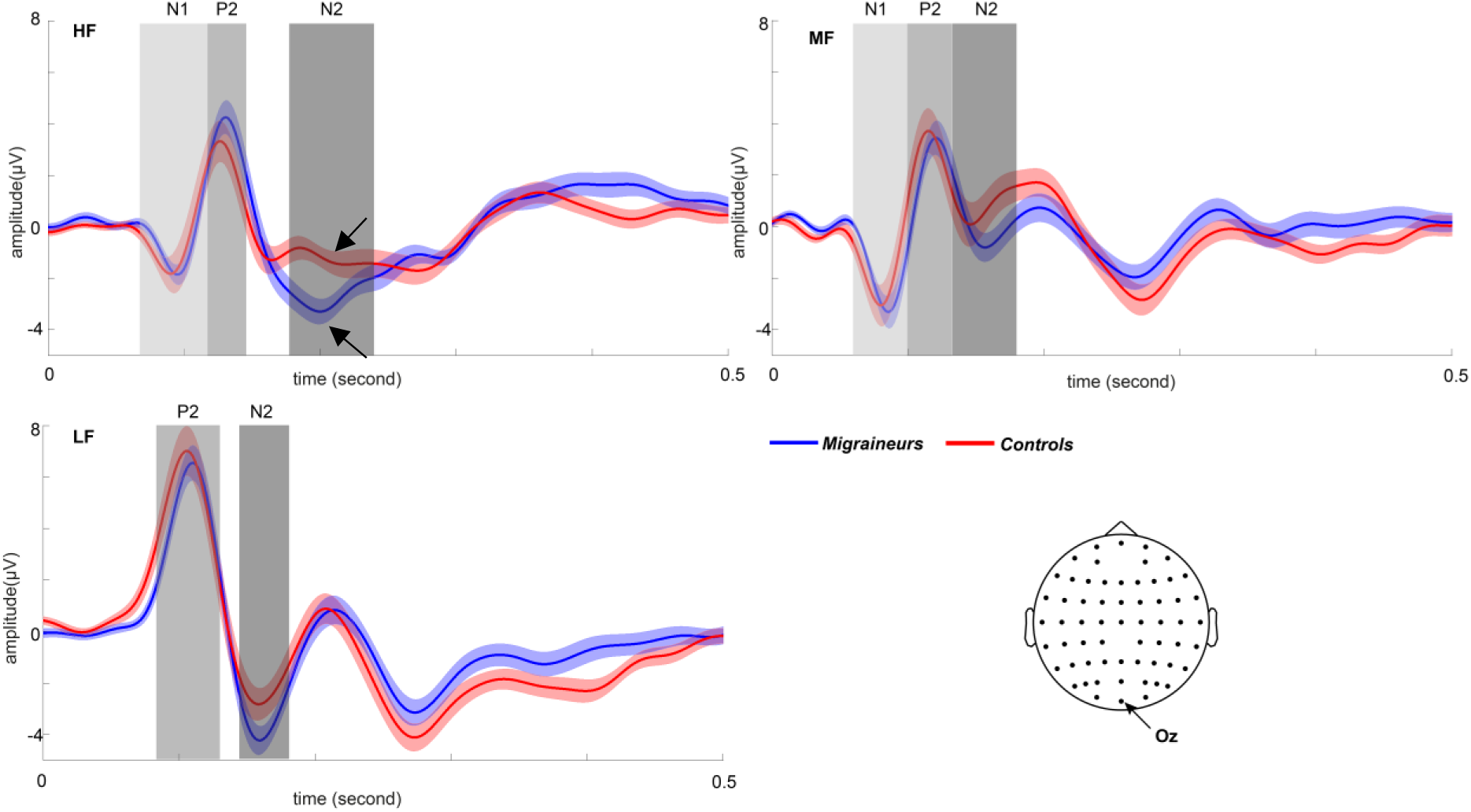
Grand mean of the ERP measured at Oz in HF (left), MF (middle) and LF (right) conditions for migraineurs vs healthy control (shaded area indicating +/-1 SE). The arrows indicated significant amplitude differences of N2 between the two groups. The time-intervals of interest used for the average peak amplitude is shaded in grey.

For exploratory purposes, we also analysed the group-differences of other early visible local peaks. The N1 was defined as the mean amplitude within the latency range of 75 –115 ms and 75 – 100 ms for HF and MF, respectively. The P2 was defined as the mean amplitude within the range 115 – 140 ms, 100 – 130 ms, 95 – 125 ms for HF, MF and LF (see Figure 4). In addition, the peak latency for each condition was measured by the local peak latency within the range defined above. However, we did not observe any other significant differences in the mean peak amplitude of the other ERP components between the two groups across all visual contrasts, all *p* > .30. Since there was no systematic latency shift between the two groups of subjects across three conditions, statistics on peak latency were not reported.

#### 3.1.2. Cluster-based Permutation Analyses

The non-parametric cluster-based permutation analysis was carried out on the 0 – 700 ms time window after the stimulus onset for three different spatial frequency conditions. The clusters were formed by significant t-stats of potential differences between migraine and control group. Given the scalp topography of the early VEP components are typically bipolar with a frontal and occipital scalp distribution, we have chosen to display only the wave-forms in significant cluster of electrodes over occipital regions.

We found that for the HF gratings, non–migraineurs relative to the migraineurs group had a significantly more negative VEP at 382 – 538 ms post-stimulus (*p* = .002, Monte Carlo *p*-value, corrected for multiple comparisons) maximally distributed over the parietal and occipital regions (see Figure 5). A similar pattern was observed over the posterior electrodes for MF (384 – 486 ms, *p* = .023, Monte Carlo *p*-value, corrected for multiple comparisons) and LF (368 – 486 ms, *p* = .012, Monte Carlo *p*-value, corrected for multiple comparisons) conditions (see Figure 6 & 7).

**Figure 5.**
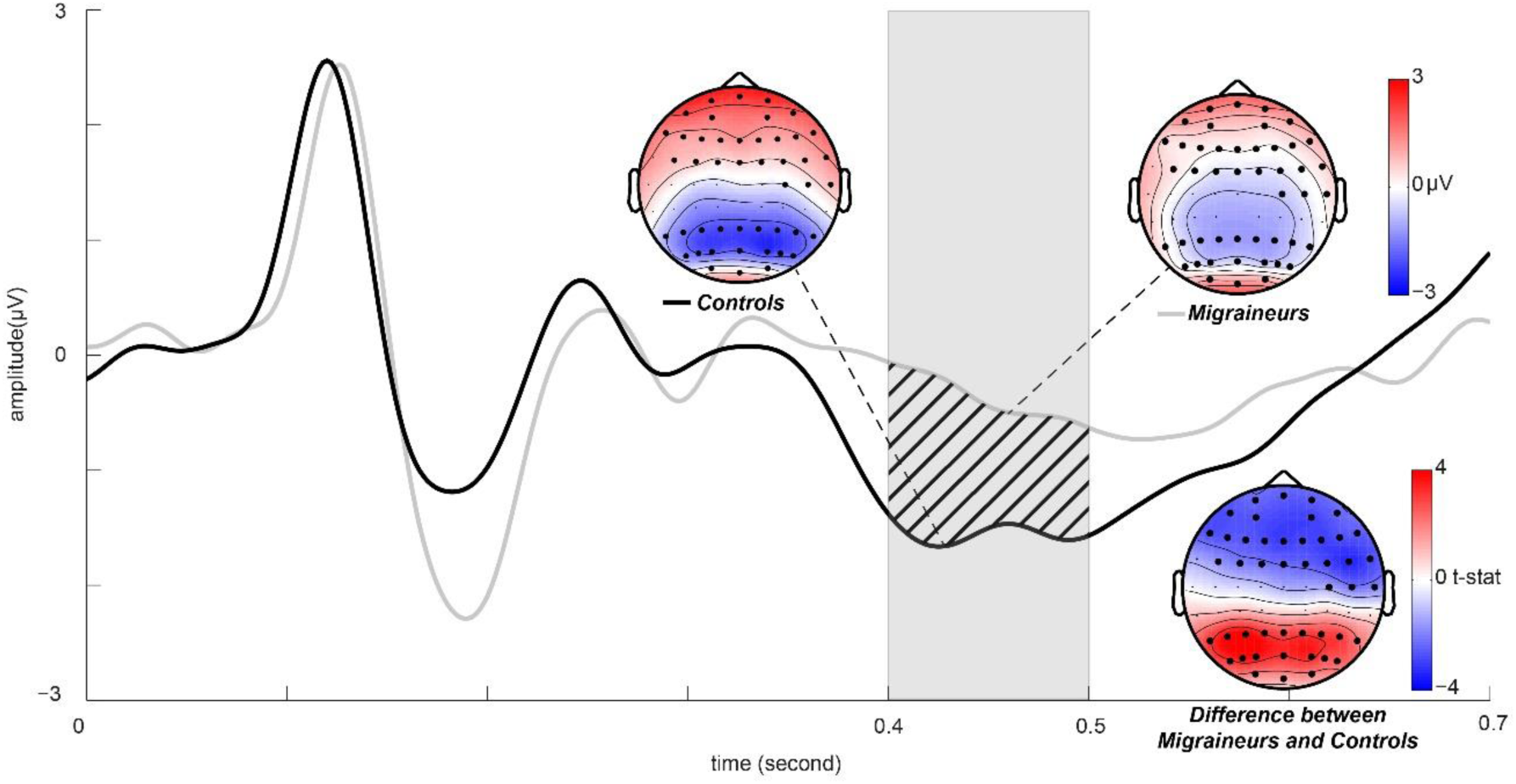
The average ERP (HF) over the significant channels of the positive cluster (posterior region)The significant channels were highlighted in bold on the topographies (both the positive and negative clusters). Migraineurs had an attenuated VEP compared to control between 400 and 500 ms (shaded in grey).

**Figure 6.**
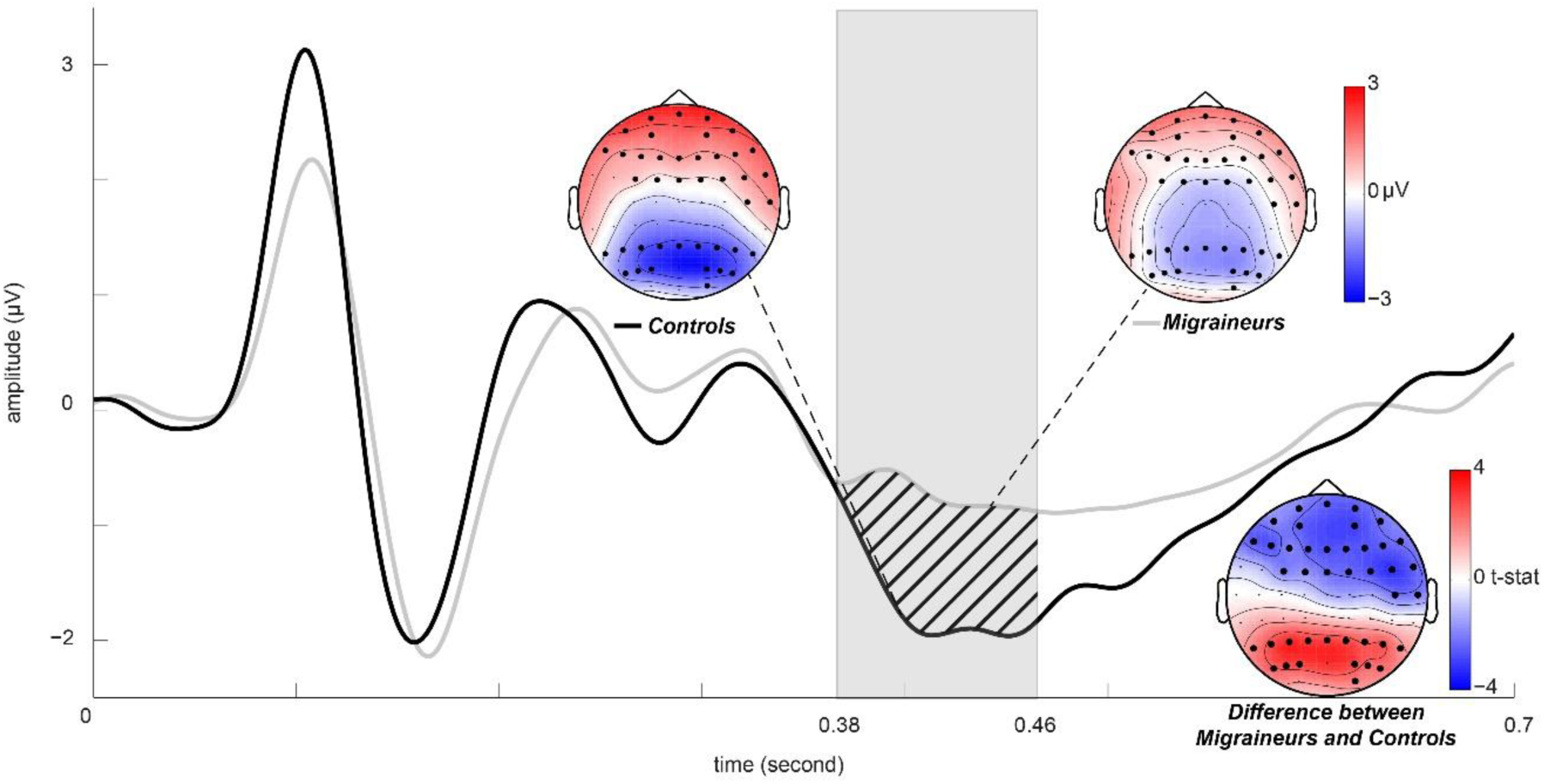
The average ERP (MF) over the significant channels of the positive cluster (posterior region).The significant channels were highlighted in bold on the topographies (both the positive and negative clusters). Migraineurs had an attenuated VEP compared to control between 380 and 460 ms (shaded in grey).

**Figure 7.**
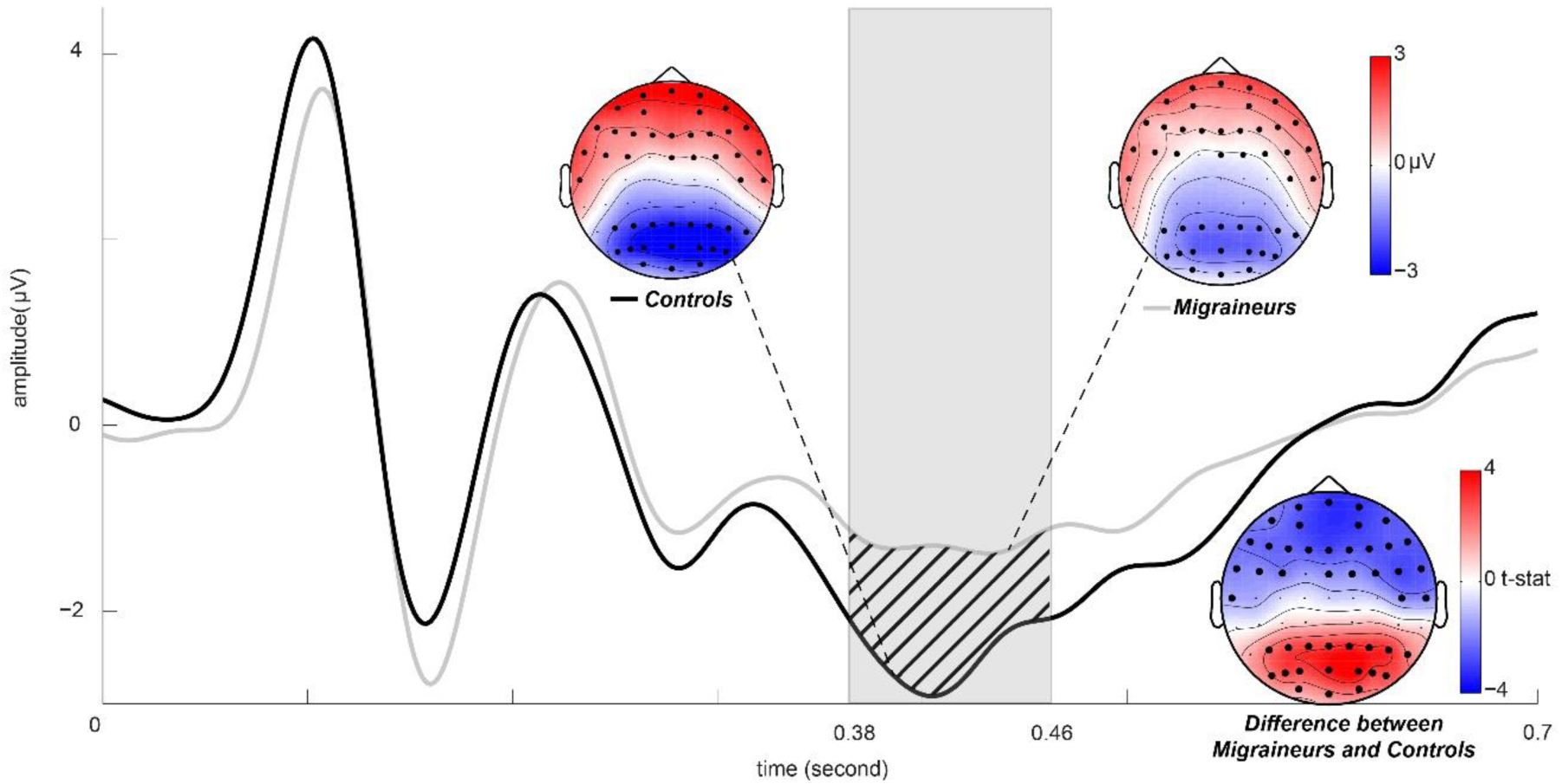
The average ERP (LF) over the significant channels of the positive cluster (posterior region). The significant channels were highlighted in bold on the topographies (both the positive and negative clusters). Migraineurs had an attenuated VEP compared to control between 380 and 460 ms (shaded in grey).

#### 3.1.3. Behavioural Tests

##### 3.1.3.1. Pattern Glare Results

Migraineurs significantly showed more AVD response at all spatial frequency conditions than control group after FDR correction (HF: *p* = 0.004; MF: *p* = 0.021; LF: *p* = 0.004; see Table 2). However, the subtraction parameter – ΔAVD (MF - HF) comparison between migraineurs and control was not significant.

**Table 2.** Mean AVD and Bayes factor for migraine vs control across HF, MF and LF conditions (with standard error in parenthesis)

| Mean AVD | Migraineurs | Control | $t$-stat | Uncorrected<br>$p$-value | FDR-corrected<br>$p$-value | $BF_{10}$ |
| --- | --- | --- | --- | --- | --- | --- |
| **HF** | **19.2 (1.94)** | **11.3 (1.56)** | **-3.20** | **.002** | **.004** | **15.8** |
| **MF** | **18.4 (2.02)** | **12.4 (1.37)** | **-3.47** | **.016** | **.021** | **3.23** |
| **LF** | **8.23 (1.29)** | **3.24 (0.70)** | **-2.48** | **.001** | **.004** | **31.2** |
| MF – HF<br>(ΔAVD) | -0.81 (1.28) | 1.14 (0.81) | 1.31 | .20 | .20 | 0.54 |

##### 3.1.3.2. CHi-II and CAPS

Migraineurs scored significantly higher than controls in *HVSD* (FDR corrected *p* < 0.001) and *AHE* (FDR corrected *p* = 0.003). Although there were mean group differences in *DVP* and *TLE*, they were marginally non-significant (see Table 3 for the mean score, *p*-value for t-test, FDR corrected *p*-value and Bayes factor).

**Table 3.** The mean questionnaire score for the subscales of *CHi-II* and *CAPS* (with standard error in parenthesis)

| Questionnaire score | Migraineurs (n = 29) | Control (n = 31) | $t$-stat | Uncorrected $p$-value | FDR corrected $p$-value | $BF_{10}$ |
| --- | --- | --- | --- | --- | --- | --- |
| ***HVSD*** | **61.1(3.74)** | **31.7(3.53)** | **-5.72** | **<.001** | **<0.001** | **>30000** |
| ***AHE*** | **24.8(3.36)** | **11.4(2.28)** | **-3.34** | **.001** | **.003** | **22.4** |
| *DVP* | 10.1(1.43) | 5.97(1.45) | -2.04 | .046 | .069 | 1.47 |
| *TLE* | 24.0(3.21) | 15.3(2.60) | -2.14 | .037 | .069 | 1.72 |
| *CS* | 12.8(2.81) | 13.3(3.34) | 0.11 | .910 | .910 | 0.26 |
| *CP* | 5.48(1.37) | 4.32(1.72) | -0.52 | .603 | .724 | 0.30 |

*Note: HVSD: Heightened Visual Sensitivity and Discomfort; AHE: Aura-Like Hallucinatory Experiences; DVP: Distorted Visual Perception; TLE: Temporal Lobe Experience; CS: Chemosensation; CP: Clinical Psychosis*

#### 3.1.4. Post-hoc Comparison for migraineurs with and without aura

We did a within a group analysis comparing the VEP of migraineurs with and without an aura. We found that the mean amplitude of N1 induced by MF gratings for the migraineurs with an aura was significantly reduced compared to the migraineurs without an aura (mean: - 1.88 vs −4.20 μV), t (27) = 2.32, *p* = .034, *BF*_*10*_ = 2.09. Such a difference between conditions was not observed in other spatial frequencies (see Figure 8). Apart from this, there were no other significant differences between the migraineurs with and without an aura in all other ERP and behavioural measures.

**Figure 8.**
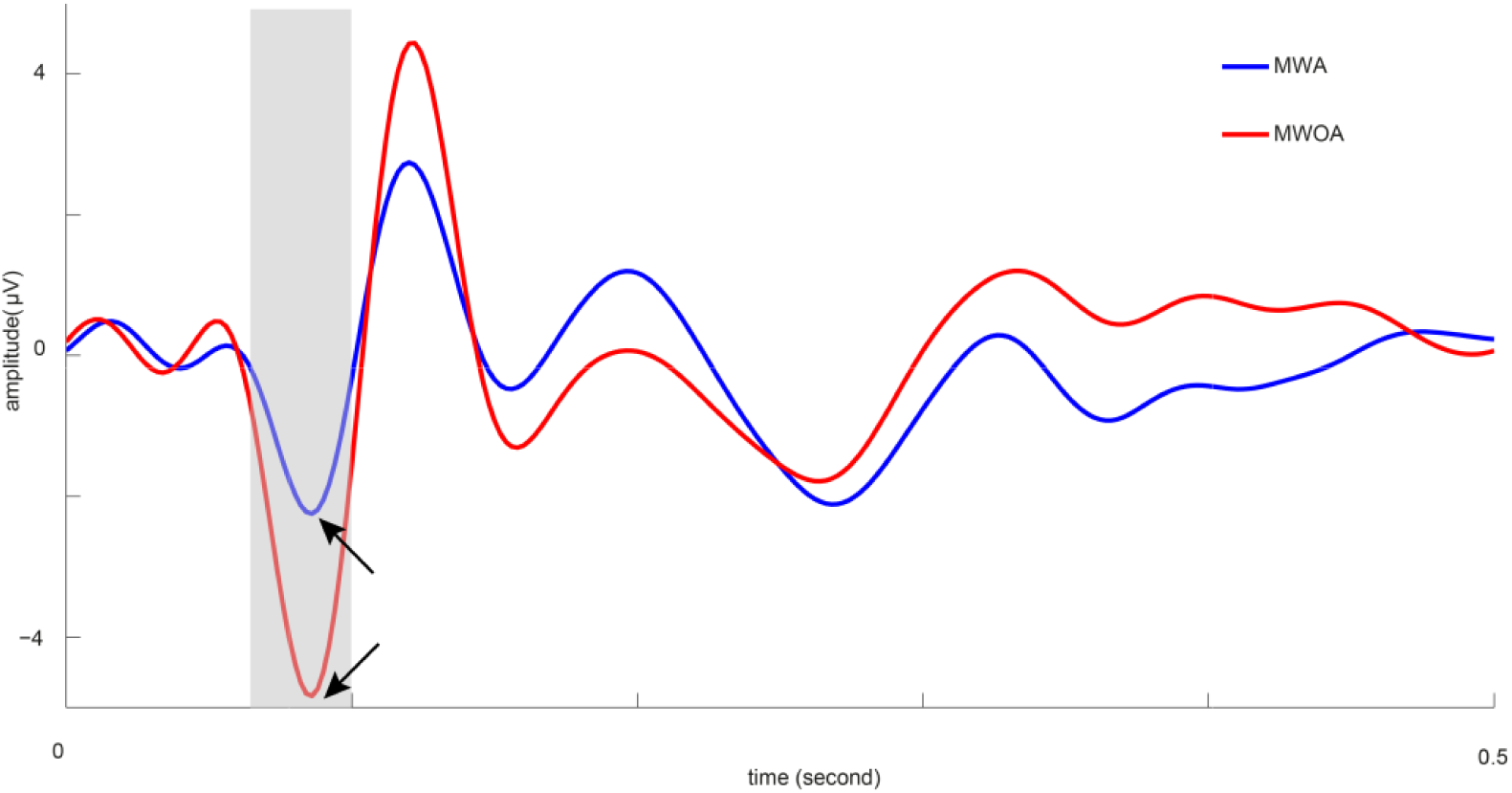
Grand mean of the ERP (MF) at Oz between migraine with aura (MWA) and migraine without aura (MWOA). There was a significant amplitude difference for N1.

### 3.2. Pattern glare (PG) vs non-Pattern glare (non-PG)

Finally, the non-migraine participants were split into two groups according to their ΔAVD score (AVD of MF subtracted by HF). The participants who have a ΔAVD > 3.92 were categorised as PG group (n = 15, mean age = 18.8) while those scoring less than 3.92 were categorised as a non-PG group (n = 23, mean age = 19.7). This reference score was obtained by a previous study (Fong et al., 2019).

As we did in the last section, the N2 component was first visually identified and defined by a latency range (HF: 150 – 180ms; MF: 130 – 165ms; LF: 140 – 180ms). The amplitude for the component was calculated by the average potential within the above time window. We found that the PG group exhibited a more negative N2 amplitude in HF condition than non-PG group significantly (mean: −2.22 μV vs −0.38 μV), t (36) = 2.176, *p* = .036, *BF*_*10*_ = 1.93 (see Figure 9; also see Figure 10 for a comparison with migraineurs).

**Figure 9.**
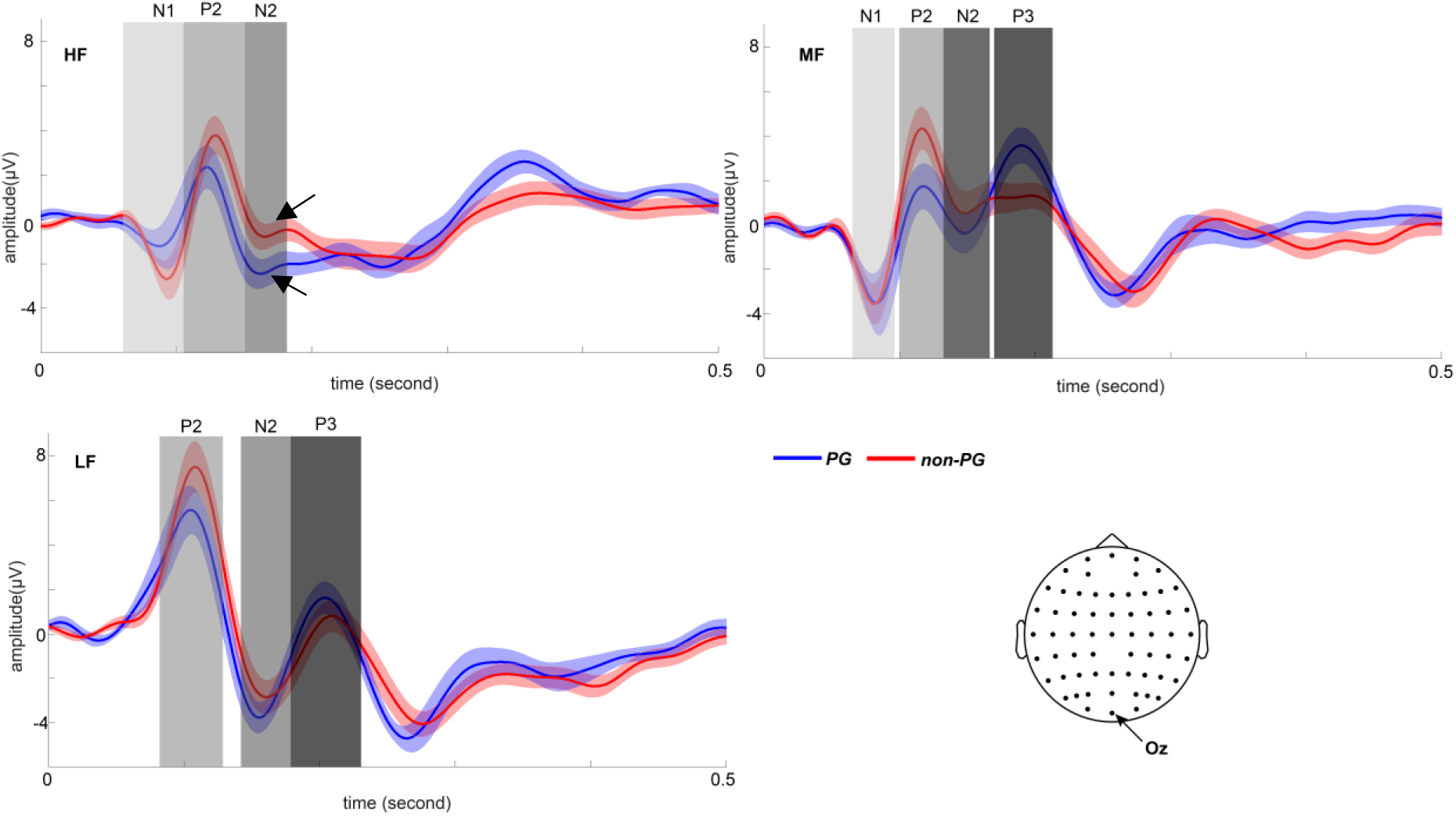
Grand mean of the ERP measured at Oz in HF, MF and LF conditions for PG group vs non-PG group (shaded area indicating +/-1 *S.E.*). The arrows indicated significant amplitude differences of N2 between the two groups. The time-intervals of interest used for the average peak amplitude is shaded in grey.

**Figure 10.**
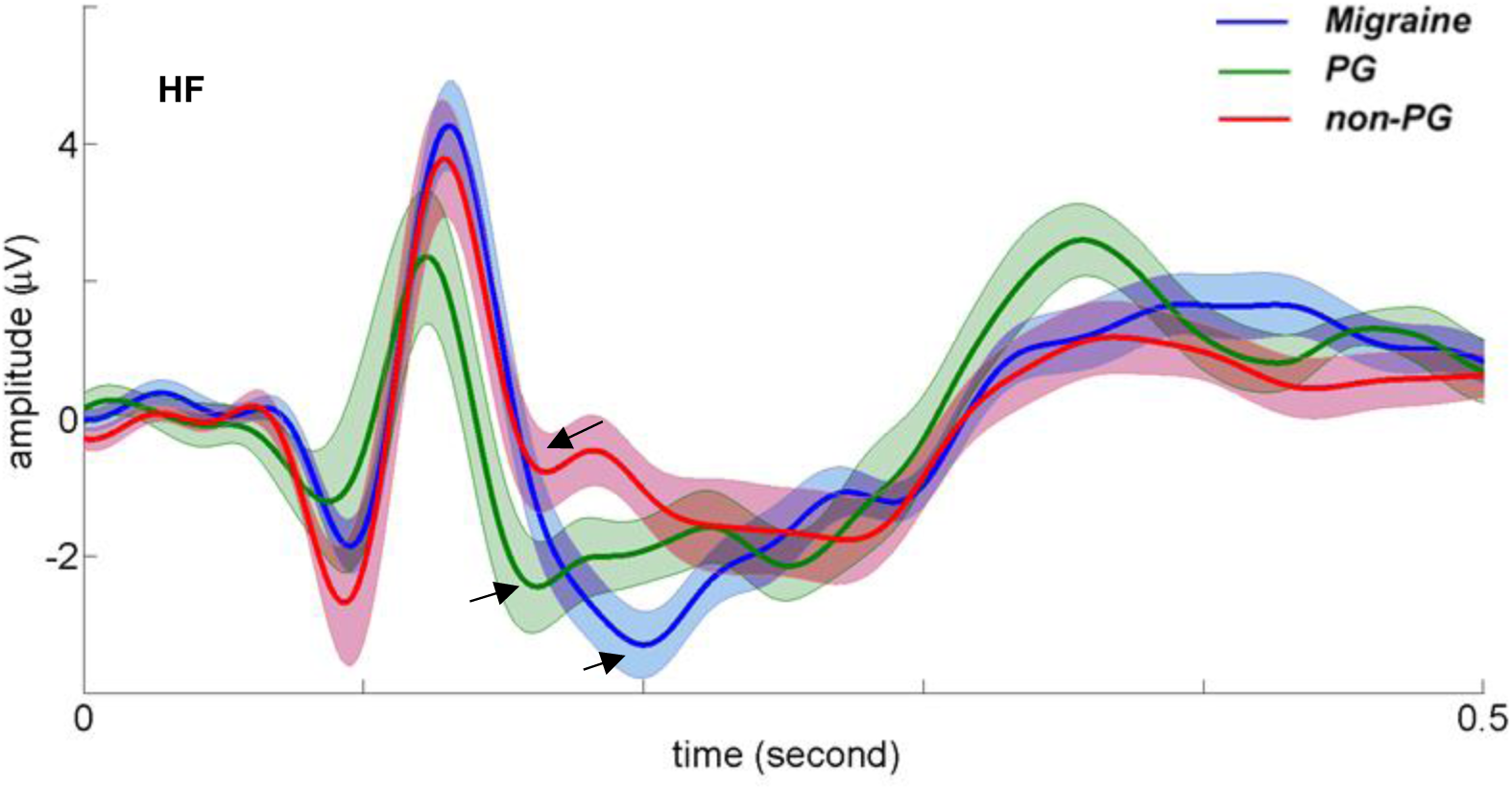
Grand mean of the ERP measured at Oz in HF, for Migraine vs PG group vs non-PG group (shaded area indicating +/-1 *S.E.*). The arrows indicated the N2 peak.

Next, any amplitude or latency differences for other VEP components (on Oz) was further explored. The latency range averaged for a peak component was shown in Table 4. In addition, latency for the components was measured as the local peak/trough within the defined latency range.

**Table 4.**
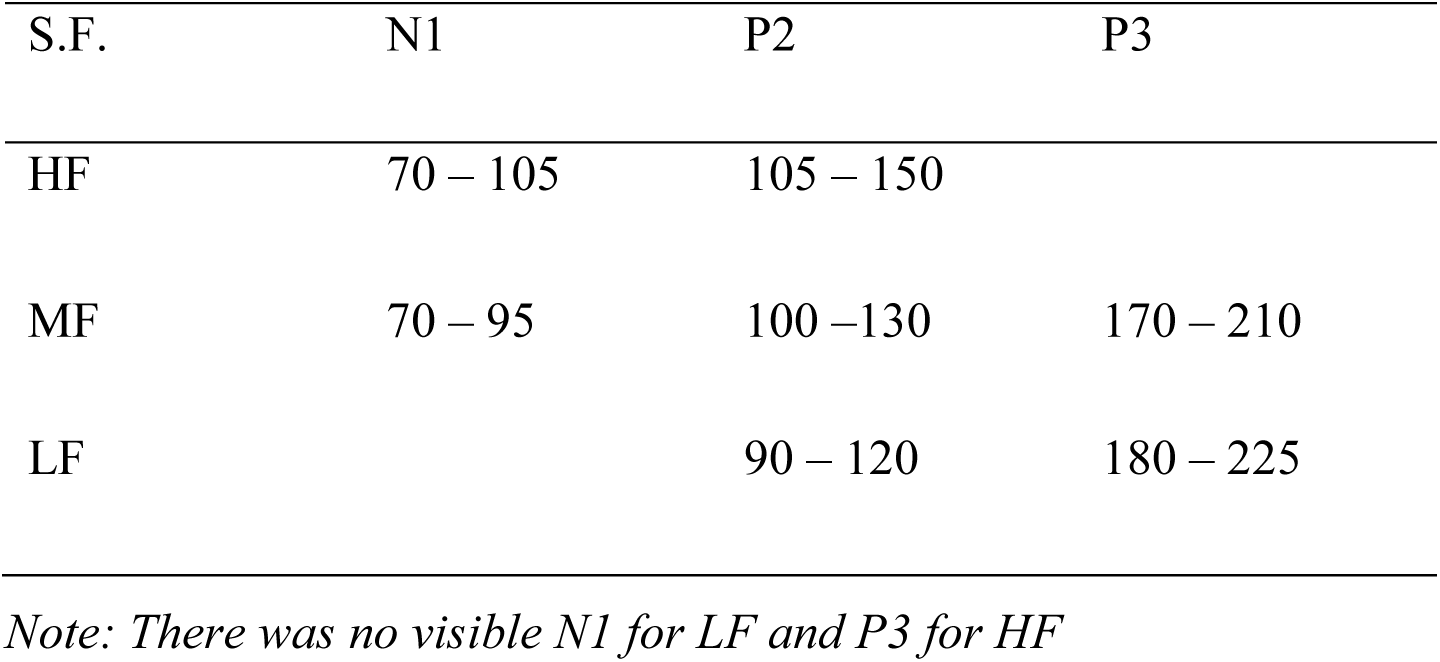
The latency range (ms) of the early VEP component in different spatial frequency (S.F.)

Our exploratory analyses discovered an increased P3 amplitude for the PG-group compared to non-PG group (mean: 2.96 μV vs 0.99 μV), t (36) = 2.213, *p* = .033, *BF*_*10*_ = 2.04 although it did not reach significance after FDR correction. These results were summarized in Table 5. Since there was no systematic latency shift between the two groups of subjects across three conditions, statistics on peak latency were not reported.

**Table 5.** Results of independent t-tests and Bayes factor on N1, P1, and P2 amplitudes between PG group and non-PG group (with *S.E.* in parenthesis).

| Mean amplitude of components (μV) | PG (n = 15) | Non-PG (n = 23) | t-stat | Uncorrected p-value | FDR corrected p-value | $BF_{10}$ |
| --- | --- | --- | --- | --- | --- | --- |
| N1 (HF) | -0.84 (0.78) | -1.28 (0.51) | -0.75 | .46 | .54 | 0.40 |
| N1 (MF) | -2.94 (1.20) | -2.26 (0.73) | 0.52 | .61 | .61 | 0.36 |
| P2 (HF) | 0.88 (0.74) | 2.42 (0.66) | 1.52 | .14 | .28 | 0.78 |
| P2 (MF) | 1.08 (0.92) | 3.63 (0.82) | 2.02 | .05 | .18 | 1.54 |
| P2 (LF) | 4.88 (0.96) | 6.96 (0.98) | 1.44 | .16 | .28 | 0.72 |
| **P3 (MF)** | **2.96 (0.75)** | **0.99 (0.53)** | **-2.21** | **.03** | **.18** | **2.04** |
| P3 (LF) | 0.83 (0.67) | -0.34 (0.58) | -1.31 | .20 | .28 | 0.62 |

#### 3.2.2 Late components

The cluster-based permutation analyses at 300 - 700 ms revealed two marginally significant clusters in the MF conditions. The positive cluster (*p* = 0.061) involved 14 centro-parietal channels (CP2, CP6, P3, Pz, P4, CP4, P5, P1, P2, P6, PO5, PO3, PO4, PO6) between 410 – 478 ms (see Figure 11). Once again, due to the dipolar nature of the VEP topography we have chosen to show the waveforms over significant electrodes over the occipital cortex. Such clusters were not observed in the other two spatial frequencies (all *p* > 0.3).

**Figure 11.**
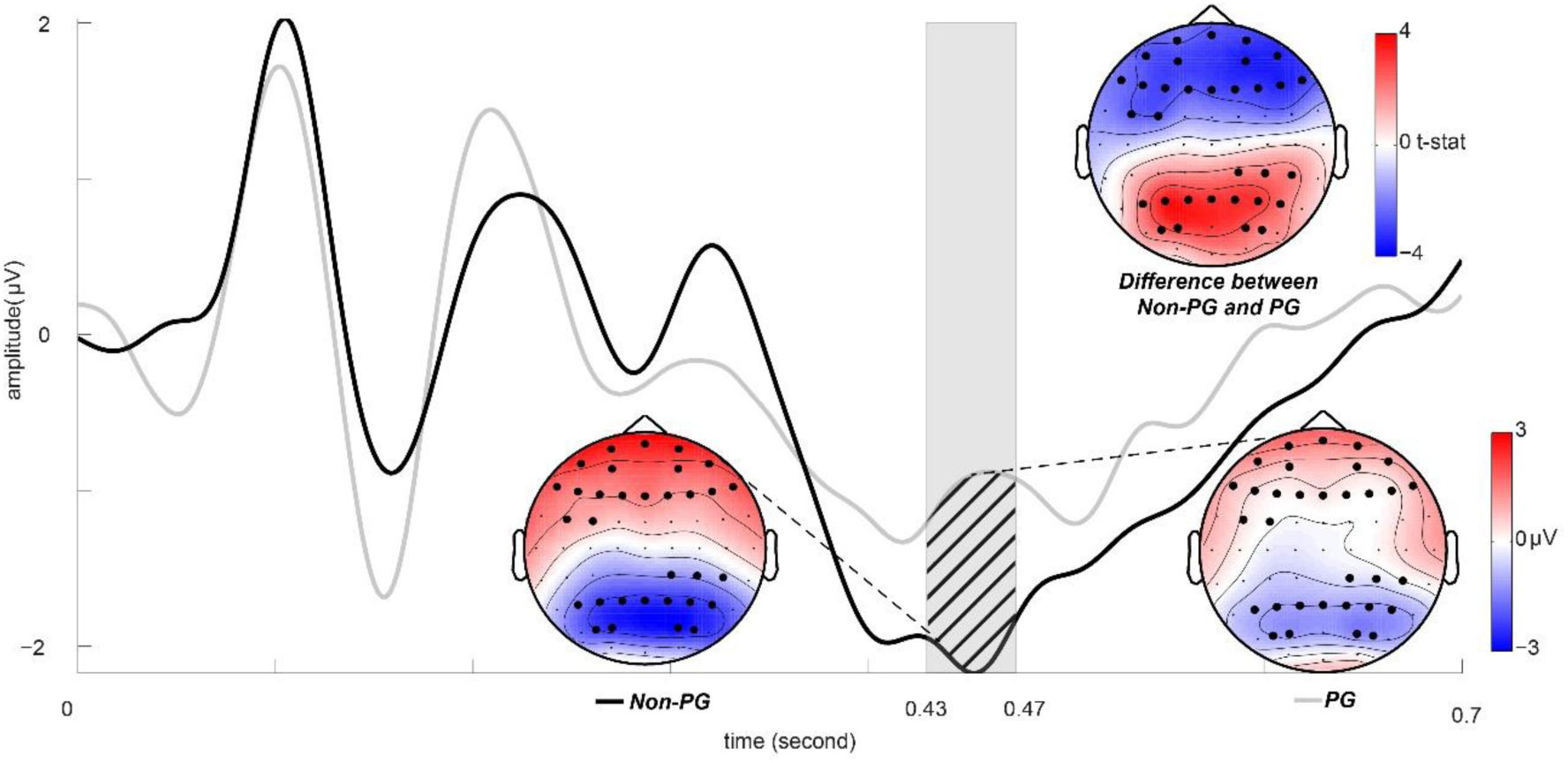
The average ERP (MF) over the marginally significant channels of the positive cluster (posterior region).The marginally significant channels were highlighted in bold on the topographies (both the positive and negative clusters). PG group had an attenuated VEP compared to non-PG group between 430 and 470 ms (shaded in grey).

#### 3.2.3. Behavioural measures – PG task and questionnaires

Behavioural measure differences between PG group and non-PG group were also explored. Table 6 showed that ΔAVD difference mainly resulted from a high mean AVD in MF for the PG group. For the questionnaire measures, there were positive correlations between the N1 amplitude (for both HF and MF gratings) and the *HVSD* score (Spearman’s rho: N1(HF) vs *HVSD* = .330, *p* = .040; N1(MF) vs *HVSD* = .346, *p* = .033), meaning a higher *HVSD* score was associated with a less negative/reduced N2 amplitude.

**Table 6.** Mean AVD for PG and non-PG across HF, MF and LF conditions (with *S.E.* in parenthesis)

| Mean AVD | PG | Non-PG |
| --- | --- | --- |
| HF | 12.4 (1.94) | 12.2 (1.56) |
| **MF** | **16.4 (2.02)** | **11.5 (1.37)** |
| LF | 4.87 (1.29) | 3.52 (0.70) |

## 4. Discussion

In the current study, we used both a confirmatory and exploratory approach to compare the in VEPs elicited by gratings of different spatial frequencies between a group of migraineurs and non-migraineurs. We found that after the presentation of the high-frequency gratings (13 cpd), the migraineurs exhibited an enhanced negative N2. A within-group analysis found that this was observed in both migraine sufferers with and without auras. However, a post-hoc analysis did reveal an N1 difference with migraine sufferers with an aura having a significantly reduced N1 amplitude than those without aura. Our exploratory analysis showed, for all gratings, an attenuation of the post-stimulus (400-500ms) LN over occipital channels in the migraine group. In the second part of the analysis, the PG group appeared to have similar VEP responses as migraineurs, with an increased N2 for HF and decreased LN for MF.

### 4.1 Findings on migraine

#### 4.1.1. Increment of N2 complex revealed cortical hyperexcitability

The enhanced N2 we observed is in line with previous studies using pattern reversal VEP paradigm (Lahat, Nadit, & Barr, et al., 1997; Lahat, Barr, & Barzilai et al., 1999; Shibata et al., 1997). Oelkers et al. (1999) proposed that the N2 complex in medium (or high) frequency grating could be a superposition of N130 and N180 components. N130 is described as contour specific and visible for both migraine and healthy subjects evoked by high-frequency grating while N180 is luminance dependent and relatively predominating in the N2 complex for migraineurs (Oelkers et al., 1999). Consistent with Oelkers et al. (1999)’s findings, the N2 in the present study for migraineurs peaks at 200 ms instead of the commonly reported 130 – 145 ms suggested that it was predominated by a luminance-dependent N180. It is, however, not clear whether there is any amplitude change for N130. Nonetheless, the abnormal N2 could reflect an imbalance between the magnocellular (sensitive to luminance: N180) and parvocellular (sensitive to contrast and high spatial frequency: N130) system. The impairment in the connectivity between these two systems could be potentially caused by cortical hyperexcitation or abnormality from GABAergic inhibitory interneurons (Chronicle and Mulleners, 1994).

However, one might argue that if the increased N2 is driven by cortical hyperexcitability, a similar N2 deflection should be observed on MF condition as well. One explanation for this divergence could be that the N2 components increased with the spatial frequency of the visual input and as such only becomes visible for high-frequency condition. This is consistent with previous literature showing an enhancement of the early negative potentials as a function of grating frequency (Oeklers et al, 1999; Hudnell et al, 1990). An alternative but not mutually exclusive explanation is that there is a “phantom” positive component, namely P200 cancelled out the negativity of N180. Some literatures showed that visual P200 is associated with motion onset (Schulte-Korne, Deimel, & Remschmidt, 2004). Although the presentation of the stimuli was steady in the experiment, the 3 cpd grating could cause a spread of discharge beyond V1 to motion perception related regions such as V3 and V5, leading to illusions of movement (Evans & Stevenson, 2008; Ffytche, Skidmore, & Zeki, 1995). This mechanism could also explain why jittering and shimming are so common as a form of motion illusions induced by gratings in 3 cpd (see Braithwaite et al., 2015; Evans & Stevenson, 2008; Fong at al., 2019). The hypothesised VEP model and the role of N130, N180 and the “phantom” P200 were summarised in Figure 12.

**Figure 12.**
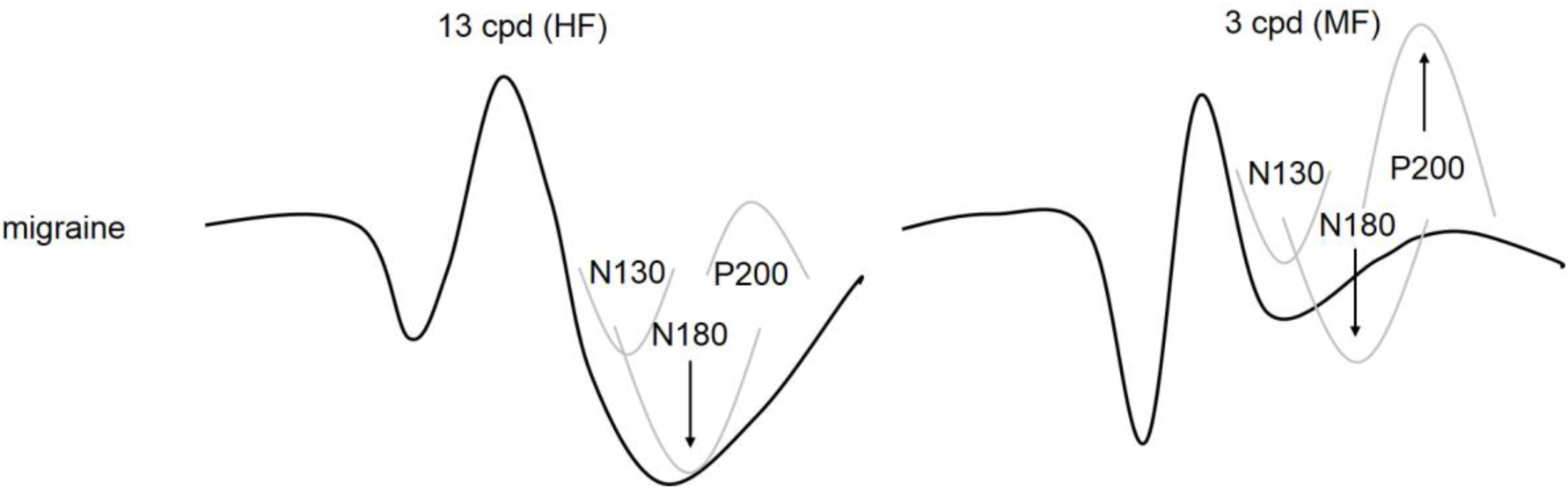
Model of the migraine VEPs for HF and MF. The N2 complex can be hypothesized as a superposition of N130, N180 and P200. For the HF condition, migraineurs had an increased luminance-dependent N180. The predominating N180 outweighed the P200. For the MF condition, the N180 increase was cancelled out by a sharp P200.

#### 4.1.2. Pathological difference between migraine subtypes

Another finding related to sensory processing is that our migraine with aura group had a significantly attenuated N1 amplitude than migraine without aura group for the MF condition, in line with previous studies (see Coppola et al., 2015; Khalil, Legg, & Anderson, 2000). The amplitude difference in early VEP components between the migraine subtypes is said to be linked with the presence and history of aura, as discovered by above literatures who report similar findings. They suggested that the presence and a prolonged suffer from aura in the long run (history) will reduce the amplitudes in early components, probably through ischaemia-induced neural damage during the experience of aura. However, it is unlikely that the N1 difference in the present study was due to neural damage from aura history with a young age sample.

On the other hand, we could argue that the present finding was due to enhanced cortical hyperexcitability between migraine sufferers with and without an aura. Although both migraine sufferers (with/without aura) were known to have elevated cortical hyperexcitability, a different dimension of cortical hyperexcitability could underlie their pathological differences. In other words, migraineurs with and without aura share the same elevated cortical hyperexcitability on one dimension but differ on another dimension. This multi-dimensional concept of cortical hyperexcitability was proposed by our previous study (Fong et al., 2019).

A study had demonstrated that later VEP components could be enhanced by altering cortical hyperexcitability through rTMS while N1 and P1 remained unchanged after receiving the same stimulation (Thut, Theoret, Pfenning, Ives, & Kampmann, 2003). Later, Di Russo et al. (2005)’s VEP-fMRI study confirmed that the visual N1 and the later component N2 might be originated from a different neural generator. These literatures appeared to support the multi-dimension model of cortical hyperexcitability and provide an explanation to the pathological difference between migraine subgroups.

#### 4.1.3. Late stage visual processing on gratings

Unlike the trend in early VEP components, there were no significant differences between the two migraine subgroups in late ERPs according to the cluster-based permutation analysis. However, significant group differences in late ERP amplitudes between migraine patients and healthy controls were obtained across all spatial frequencies. These differences were denoted by the LN (centred at parietal and occipital-temporal areas) peaked around 400 – 500 ms. Such activities were significantly attenuated in the migraine sample, i.e. a reduction in amplitude in LN was observed.

Late potentials (LP) are widely agreed to be associated with stimulus recognition and selective attentional processing (Cuthbert et al., 2000; Schupp, Junghofer, Weike, & Hamm, 2004; Ritter & Ruchkin, 1992). For example, affective images were known to elicit an enlarged LP compared to neutral images (Schupp et al., 2004). Migraineurs were found to have a reduced LP amplitude when affective images were presented regardless of the valance of pictures (Tommaso et al., 2009). Abnormal LP was also reported in other literatures with varied findings in uncertain directions - an increment or reduction (Mickleborough et al., 2013; 2014; Steppacher et al., 2016). Although late potentials have rarely been studied in a pVEP paradigm, it is possible that the aversive gratings induce a similar top-down bias on visual processing, leading to such an LP group difference. This explanation is also supported by our behavioural data, which has shown that migraineurs have an increased visual sensitivity at all spatial frequencies in the PG task, in line with previous research with other behavioural or physiological measures (Oelkers et al., 1999; Huang, Cooper, Satana, Kaufman, & Cao, 2003). Therefore, this top-down bias could cause visual attention inhibition and counterbalance the discomfort caused by the hypersensitivity of migraineurs on the gratings. However, we cannot rule out the possibility that our migraine sample had a general visual attention deficit regardless the context of the stimuli which were also found in the literature (Ince, Erdogan-Bakar, & Unal-Cevik, 2017; Moutran et al., 2011; Villa et al., 2009). In future studies, an appropriate baseline image, such as a non-striped pattern picture, would benefit the research by revealing whether the group effects were indeed associated with the spatial frequency of the striped patterns.

### 4.2. Findings on healthy PG group

#### 4.2.1. Evidence of Cortical Hyperexcitability Supported by Early VEP Components

The PG group showed increased N2 amplitude compared to the non-PG group for the HF grating. Interestingly, this finding was similar to the above observation in which migraineurs showed abnormal N2 response compared to healthy controls. However, the current N2 component peaked at around 150 ms instead of 200 ms, suggesting a potential difference in the underlying neurocognition between the two negative components. Based on the VEP waveform, the current group difference was more likely to be caused by the increment of N130 in the absence of a migraine-specific N180 (see Figure 13). The amplitude of N130 appeared to be increased with the spatial frequency of the grating. As mentioned above, N130 is contour-specific and therefore, provides the support that the PG group might have abnormal responses along the parvocellular pathway similar to migraine patients (Oelkers et al., 1999). Such abnormality was believed to be caused by impaired GABAergic inhibitory system, which can manifest itself as cortical hyperexcitation (Chronicle and Mulleners, 1994).

**Figure 13.**
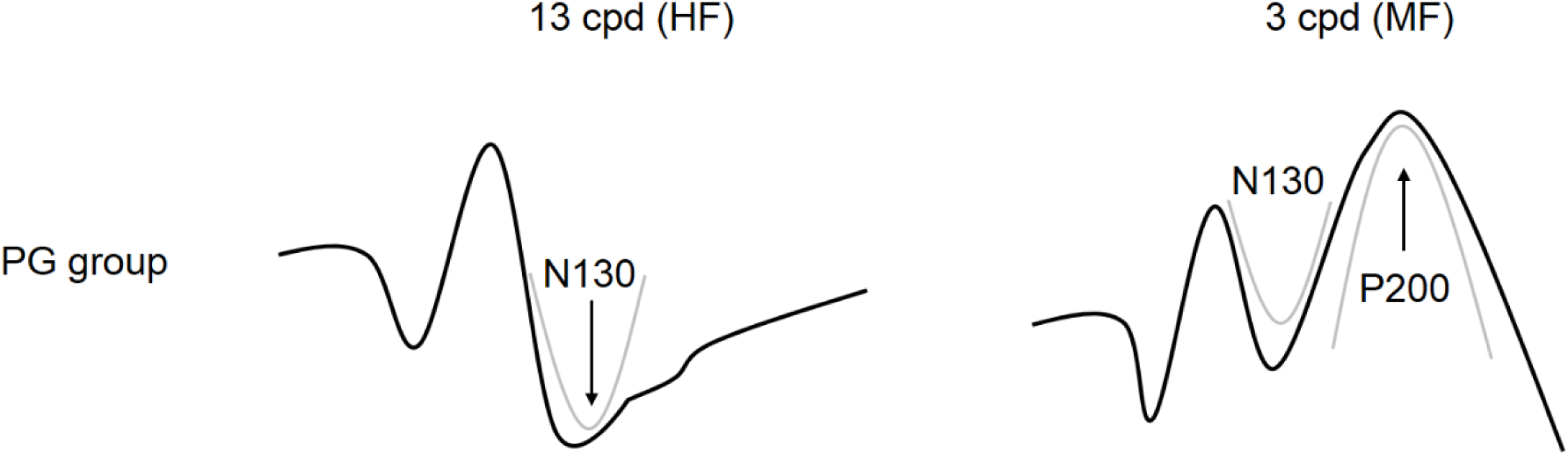
Model of the PG group VEPs for HF and MF. The N2 complex was hypothesised as a superposition of N130, N180 and P200 in the previous chapter while N180 is absent or attenuated for the present non-migraine sample. For the HF condition, PG group had an increased N130. For the MF condition, in the absence of the migraine-specific N180, the aberrant experience-associated P200 become visible.

We hypothesised that a “phantom” P200 component was cancelled out by a migraine-specific predominating N180. This P200 is also thought to be associated with the aberrant experience induced by the grating. In the sample of healthy population, with the absence of N180, an increased P200 (shown as P3 in Figure 9) was observed on the PG group (who experienced excessive pattern glare in MF grating) compared to the non-PG group. The model of VEP for MF was summarised in Figure 13.

Collectively, our findings provide further support that cortical hyperexcitability could be the basis of the experienced pattern-glare effect but now further extended to non-clinical groups and to specific EEG components.

The N1 for MF and HF were found to be positively correlated (Spearman’s rho = .346 & .330) with one of the *CHi-II* factors – namely *HVSD.* If the neural source of this N1 component was the same as the one highlighting the migraine subgroup difference (e.g. V1), then this dimension of cortical hyperexcitability could underlie everyday life pattern or light-induced visual stress symptoms as well as the pathological difference of migraine subtype.

#### 4.2.2. Evidence of Selective Visual Processing by Late VEP

Apart from sensory processing, VEP components peaked at around 200 – 300 ms can also reflect higher-order cognitive modulation such as selective attention on spatial frequency (Proverbio, Esposito, & Zani, 2002; Zani & Proverbio, 1995, 2009). For instance, P200 was believed to be responsible for selective attention (Hackley, Woldorff, & Hillyard, 1990; Noldy, Stelmack, & Campbell, 1990) and features detection (Luck & Hillyard, 1994). In our current setting, we are unable to conclude the source of the P3 deflection on the hyperexcitable participants. However, such deflection was not observed in other spatial frequencies which is consistent with the selective attention theory (Zani & Proverbio, 2009). Whether this finding implies an attentional enhancement of the PG group on MF due to their illusive perception would require more in-depth investigation.

Though the cluster-based analysis was just marginally significant, the attenuated late negativity (LN) on PG group in MF could be another supportive evidence on selective attention on spatial frequency. Based on the migraine sample, we proposed that the LN reduction could be either caused by a top-down visual attention inhibition or a general visual attention deficit. Results in this part seemed to support the former theory as the LN reduction only appeared in MF. Since subjects in PG group are more averse to MF grating, this top-down processing could counterbalance their hypersensitivity as well as the earlier attentional enhancement by disengaging from the stimuli.

However, the limitation of our interpretation is that attention was not manipulated in our experiment apart from the verbal instructions telling the subjects to concentrate on the fixation spot. Therefore, whether the effect on LN was caused by attention has to be further investigated. For instance, an unattended condition of the grating presentation could be introduced in further study. As a result, we would see a clear picture of the role of visual attention in perceiving aversive gratings.

### 4.3. Conclusion

In conclusion, migraine patients have an increased N2 amplitude in HF gratings compared to controls, hinting a sensory impairment consistent with the cortical hyperexcitability mechanism. In addition, pathological differences between migraine subgroups are supported by a significantly higher N1 amplitude in the MWOA than MWA group. Apart from the early VEP, the late negativity across all spatial frequencies differ in migraineurs and healthy controls may imply migraineurs’ attentional inhibition to the striped patterns. Some current findings on healthy PG group are consistent with results on migraineurs. The similarity highlighted the contribution of cortical hyperexcitability to pattern induced visual disturbances and top-down suppressive control on visual gratings.

## References

Afra, J., Cecchini, A. P., De Pasqua, V., Albert, A., & Schoenen, J. (1998). Visual evoked potentials during long periods of pattern-reversal stimulation in migraine. Brain: a journal of neurology, 121(2), 233–241.

Ambrosini, A., & Schoenen, J. (2006). Electrophysiological response patterns of primary sensory cortices in migraine. The journal of headache and pain, 7(6), 377.

Aurora, S. K., & Wilkinson, F. (2007). The brain is hyperexcitable in migraine. Cephalalgia, 27(12), 1442–1453.

Boćkowski, L., Sobaniec, W., Sołowiej, E., & Smigielska-Kuzia, J. (2004). Auditory cognitive event-related potentials in migraine with and without aura in children and adolescents. Neurologia i neurochirurgia polska, 38(1), 9–14.Chen et al., 2007;

Chronicle, E., & Mulleners, W. (1994). Might migraine damage the brain?. Cephalalgia, 14(6), 415–418.

Coppola, G., Bracaglia, M., Di Lenola, D., Di Lorenzo, C., Serrao, M., Parisi, V., … & Pierelli, F. (2015). Visual evoked potentials in subgroups of migraine with aura patients. The journal of headache and pain, 16(1), 92.

Cuthbert, B. N., Schupp, H. T., Bradley, M. M., Birbaumer, N., & Lang, P. J. (2000). Brain potentials in affective picture processing: covariation with autonomic arousal and affective report. Biological psychology, 52(2), 95–111.Delorme & Makeig, 2004

Di Russo, F., Pitzalis, S., Spitoni, G., Aprile, T., Patria, F., Spinelli, D., & Hillyard, S. A. (2005). Identification of the neural sources of the pattern-reversal VEP. Neuroimage, 24(3), 874–886.

Drake Jr, M. E., Pakalnis, A., & Padamadan, H. (1989). Long-latency auditory event related potentials in migraine. Headache: The Journal of Head and Face Pain, 29(4), 239–241.

Ellemberg, D., Hammarrenger, B., Lepore, F., Roy, M. S., & Guillemot, J. P. (2001). Contrast dependency of VEPs as a function of spatial frequency: the parvocellular and magnocellular contributions to human VEPs. Spatial Vision, 15(1), 99–112.

Evans, B. J. W., & Stevenson, S. J. (2008). The Pattern Glare Test: a review and determination of normative values. Ophthalmic and Physiological Optics, 28(4), 295–309.

Evans, B. J., Cook, A., Richards, I. L., & Drasdo, N. (1994). Effect of pattern glare and colored overlays on a stimulated-reading task in dyslexics and normal readers. Optometry and vision science: official publication of the American Academy of Optometry, 71(10), 619–628.

Ferree, T. C. (2000). Spline interpolation of the scalp EEG. Secondary TitlEGI.

Ffytche, D. H., Skidmore, B. D., & Zeki, S. (1995). Motion-from-hue activates area V5 of human visual cortex. Proceedings of the Royal Society of London. Series B: Biological Sciences, 260(1359), 353–358.

Fong, C. Y., Takahashi, C., & Braithwaite, J. J. (2019). Evidence for distinct clusters of diverse anomalous experiences and their selective association with signs of elevated cortical hyperexcitability. Consciousness and cognition, 71, 1–17.

Foxe, J. J., Strugstad, E. C., Sehatpour, P., Molholm, S., Pasieka, W., Schroeder, C. E., & McCourt, M. E. (2008). Parvocellular and magnocellular contributions to the initial generators of the visual evoked potential: high-density electrical mapping of the “C1” component. Brain topography, 21(1), 11–21.

Hackley, S. A., Woldorff, M., & Hillyard, S. A. (1990). Cross-modal selective attention effects on retinal, myogenic, brainstem, and cerebral evoked potentials. Psychophysiology, 27(2), 195–208.

Harle, D. E., Shepherd, A. J., & Evans, B. J. (2006). Visual stimuli are common triggers of migraine and are associated with pattern glare. Headache: The Journal of Head and Face Pain, 46(9), 1431–1440.

Hatanaka, K., Nakasato, N., Seki, K., Kanno, A., Mizoi, K., & Yoshimoto, T. (1997). Striate cortical generators of the N75, P100 and N145 components localized by pattern reversal visual evoked magnetic fields. The Tohoku journal of experimental medicine, 182(1), 9–14. Huang, Cooper, Satana, Kaufman, & Cao, 2003

Hudnell, H. K., Boyes, W. K., & Otto, D. A. (1990). Stationary pattern adaptation and the early components in human visual evoked potentials. Electroencephalography and Clinical Neurophysiology/Evoked Potentials Section, 77(3), 190–198.

Ince, F., Erdogan-Bakar, E., & Unal-Cevik, I. (2017). Preventive drugs restore visual evoked habituation and attention in migraineurs. Acta Neurologica Belgica, 117(2), 523–530.

Khalil, N. M., Legg, N. J., & Anderson, D. J. (2000). Long term decline of P100 amplitude in migraine with aura. Journal of Neurology, Neurosurgery & Psychiatry, 69(4), 507–511.

Lahat, E., Berkovitch, M., Barr, J., Paret, G., & Barzilai, A. (1999). Abnormal visual evoked potentials in children with” Alice in Wonderland” syndrome due to infectious mononucleosis. Journal of child neurology, 14(11), 732–735.

Lahat, E., Nadir, E., Barr, J., Eshel, G., Aladjem, M., & Biatrilze, T. (1997). Visual evoked potentials: a diagnostic test for migraine headache in children. Developmental Medicine & Child Neurology, 39(2), 85–87.

Lehmann, D., Darcey, T. M., & Skrandies, W. (1982). Intracerebral and scalp fields evoked by hemiretinal checkerboard reversal, and modeling of their dipole generators. Advances in neurology, 32, 41–48.

Luck, S. J., & Hillyard, S. A. (1994). Spatial filtering during visual search: evidence from human electrophysiology. Journal of Experimental Psychology: Human Perception and Performance, 20(5), 1000.

Maris, E., & Oostenveld, R. (2007). Nonparametric statistical testing of EEG-and MEG-data. Journal of neuroscience methods, 164(1), 177–190.

Matsui, H., Udaka, F., Tamura, A., Oda, M., Kubori, T., Nishinaka, K., & Kameyama, M. (2005). The relation between visual hallucinations and visual evoked potential in Parkinson disease. Clinical neuropharmacology, 28(2), 79–82.

Mazzotta, G., Alberti, A., Santucci, A., & Gallai, V. (1995). The event-related potential P300 during headache-free period and spontaneous attack in adult headache sufferers. Headache: The Journal of Head and Face Pain, 35(4), 210–215.

Mickleborough, M. J., Chapman, C. M., Toma, A. S., Chan, J. H., Truong, G., & Handy, T. C. (2013). Interictal neurocognitive processing of visual stimuli in migraine: evidence from event-related potentials. PloS one, 8(11), e80920.

Moutran, A. R. C., Villa, T. R., Diaz, L. A. S., Noffs, M. H. D. S., Pinto, M. M. P., Gabbai, A. A., & Carvalho, D. D. S. (2011). Migraine and cognition in children: a controlled study. Arquivos de neuro-psiquiatria, 69(2A), 192–195.

Noldy, N. E., Stelmack, R. M., & Campbell, K. B. (1990). Event-related potentials and recognition memory for pictures and words: The effects of intentional and incidental learning. Psychophysiology, 27(4), 417–428.

Oelkers, R., Grosser, K., Lang, E., Geisslinger, G., Kobal, G., Brune, K., & Lötsch, J. (1999). Visual evoked potentials in migraine patients: alterations depend on pattern spatial frequency. Brain, 122(6), 1147–1155.

Oostenveld, R., Fries, P., Maris, E., & Schoffelen, J. M. (2011). FieldTrip: open source software for advanced analysis of MEG, EEG, and invasive electrophysiological data. Computational intelligence and neuroscience, 2011, 1.

Picton, T. W. (1992). The P300 wave of the human event-related potential. Journal of clinical neurophysiology, 9(4), 456–479.

Proverbio, A. M., Esposito, P., & Zani, A. (2002). Early involvement of the temporal area in attentional selection of grating orientation: an ERP study. Cognitive Brain Research, 13(1), 139–151.

Puca, F., & De Tommaso, M. (1999). Clinical neurophysiology in childhood headache. Cephalalgia, 19(3), 137–146.

Rady, A., Elsheshai, A., Elkholy, O., El-Wafa, H. A., & Ramadan, I. (2011). Visual Evoked Potential (VEP) in schizophrenia and psychotic depression. World Journal of Life Sciences and Medical Research, 1(2), 11–14.

Ritter, W., & Ruchkin, D. S. (1992). A Review of Event-Related Potential Components Discovered in the Context of Studying P3 a. Annals of the New York Academy of Sciences, 658(1), 1–32.

Saitoh, E., Adachi-Usami, E., Mizota, A., & Fujimoto, N. (2001). Comparison of visual evoked potentials in patients with psychogenic visual disturbance and malingering. Journal of pediatric ophthalmology and strabismus, 38(1), 21–26.

Schroeder, C. E., Steinschneider, M., Javitt, D. C., Tenke, C. E., Givre, S. J., Mehta, A. D., … & Vaughan, J. H. (1995). Localization of ERP generators and identification of underlying neural processes. Electroencephalography and clinical neurophysiology. Supplement, 44, 55–75.

Schulte-Körne, G., Bartling, J., Deimel, W., & Remschmidt, H. (2004). Motion-onset VEPs in dyslexia. Evidence for visual perceptual deficit. Neuroreport, 15(6), 1075–1078. Thut, Theoret, Pfenning, Ives, Kampmann, 2003

Schupp, H. T., Junghöfer, M., Weike, A. I., & Hamm, A. O. (2004). The selective processing of briefly presented affective pictures: an ERP analysis. Psychophysiology, 41(3), 441–449.

Shibata, K., Osawa, M., & Iwata, M. (1997). Pattern reversal visual evoked potentials in classic and common migraine. Journal of the neurological sciences, 145(2), 177–181.

Shibata, K., Osawa, M., & Iwata, M. (1998). Pattern reversal visual evoked potentials in migraine with aura and migraine aura without headache. Cephalalgia, 18(6), 319–323.

Shibata, K., Yamane, K., Iwata, M., & Ohkawa, S. (2005). Evaluating the effects of spatial frequency on migraines by using pattern-reversal visual evoked potentials. Clinical Neurophysiology, 116(9), 2220–2227.

Souza, G. S., Gomes, B. D., Lacerda, E. M. C., Saito, C. A., Da Silva Filho, M., & Silveira, L. C. L. (2008). Amplitude of the transient visual evoked potential (tVEP) as a function of achromatic and chromatic contrast: contribution of different visual pathways. Visual neuroscience, 25(3), 317–325.

Steppacher, I., Schindler, S., & Kissler, J. (2016). Higher, faster, worse? An event-related potentials study of affective picture processing in migraine. Cephalalgia, 36(3), 249–257.

Tagliati, M., Sabbadini, M., Bernardi, G., & Silvestrini, M. (1995). Multichannel visual evoked potentials in migraine. Electroencephalography and Clinical Neurophysiology/Evoked Potentials Section, 96(1), 1–5.

Thut, G., Theoret, H., Pfennig, A., Ives, J., Kampmann, F., Northoff, G., & Pascual-Leone, A. (2003). Differential effects of low-frequency rTMS at the occipital pole on visual-induced alpha desynchronization and visual-evoked potentials. Neuroimage, 18(2), 334–347.Tommaso et al., 2009;

Vanni, S., Warnking, J., Dojat, M., Delon-Martin, C., Bullier, J., & Segebarth, C. (2004). Sequence of pattern onset responses in the human visual areas: an fMRI constrained VEP source analysis. Neuroimage, 21(3), 801–817.

Villa, T. R., Moutran, A. C., Diaz, L. S., Pinto, M. P., Carvalho, F. A., Gabbai, A. A., & de Souza Carvalho, D. (2009). Visual attention in children with migraine: a controlled comparative study. Cephalalgia, 29(6), 631–634.

Wilkins, A. J. (1995). Visual stress. Oxford University Press.

Yilmaz, M., Erdal, M. E., Herken, H., Cataloluk, O., Barlas, Ö., & Bayazıt, Y. A. (2001). Significance of serotonin transporter gene polymorphism in migraine. Journal of the neurological sciences, 186(1-2), 27–30.

Zani, A., & Proverbio, A. M. (1995). ERP signs of early selective attention effects to check size. Electroencephalography and clinical neurophysiology, 95(4), 277–292.

Zani, A., & Proverbio, A. M. (2009). Selective attention to spatial frequency gratings affects visual processing as early as 60 msec. poststimulus. Perceptual and motor skills, 109(1), 140–158.

Zani, A., & Proverbio, A. M. (2003). Cognitive electrophysiology of mind and brain. In The cognitive electrophysiology of mind and brain (pp. 3–12). Academic press.

